# Finding dugongs through environmental DNA: a complementary tool for population surveillance

**DOI:** 10.64898/2026.09.30.755531

**Authors:** Liv Aull, Christophe Cleguer, Damien Burrows, Jon Lapeyra Martin, Dean Mathews, Stephen Widdicombe, Henrik Stahl, Jan Strugnell, Charlotte Havermans, Cecilia Villacorta-Rath

## Abstract

Detecting dugongs at low densities remains a major challenge for conservation monitoring, particularly in remote or turbid coastal habitats where conventional surveys are difficult. Environmental DNA (eDNA) offers a complementary approach, but robust species-specific tools are needed before it can be confidently applied to dugong surveillance. We developed and analytically validated a quantitative PCR TaqMan assay for detecting dugong (*Dugong dugon*) eDNA and evaluated its performance across controlled aquarium conditions in Australia and Abu Dhabi, and wild populations in northern Australia and the Arabian/Persian Gulf. The assay detected dugong eDNA in aquarium and field samples, with Sanger sequencing confirming amplification of dugong mitochondrial DNA from positive field detections. Our assay provides a robust and geographically transferable tool for targeted dugong eDNA detection, performing reliably across contrasting coastal environments and widely separated populations. This approach can strengthen local-scale surveillance of low-density or difficult-to-detect dugong populations.

## 1. INTRODUCTION

Coastal tropical marine ecosystems are undergoing rapid ecological change as climate-driven disturbance, coastal development, pollution, fisheries interactions, and habitat degradation increasingly affect the species and ecological processes they support (He & Silliman 2019). Among the most vulnerable components of these systems are large, long-lived marine vertebrates that can be particularly challenging to monitor because changes in abundance or distribution may be difficult to detect without sustained, large-scale survey programs (McCauley et al. 2015; Pimiento et al. 2020).

Dugongs (*Dugong dugon*) are long-lived marine mammals distributed across tropical and subtropical coastal waters of the Indo-Pacific. As predominantly seagrass-grazing herbivores, they are strongly associated with shallow coastal habitats and can influence seagrass community structure, productivity and associated ecosystem processes through their grazing activities (Marsh et al. 2011, 2018; Wirsing et al. 2022). Dugongs are globally listed as Vulnerable on the IUCN Red List, including those in Japan and Eastern Africa, assessed as Critically Endangered, and New Caledonia, assessed as Endangered (Marsh et al. 2025b).

Australia and the Arabian/Persian Gulf support two of the world’s major remaining dugong strongholds (Marsh et al. 2025a,c). However, even within these strongholds, information on population status and change is spatially and temporally uneven. Across most of their range in Australia, population surveys are infrequently conducted. The most frequently surveyed populations are along the coast of the Great Barrier Reef (Australia), where repeated aerial surveys have revealed a long-term decline in dugong abundance between the southerly half of the GBR, between Mission Beach and Bundaberg. The most recent survey in 2022 estimated 2,124 ± SE 476 dugongs and identified an estimated rate of decline of 2.3% per year between 2005 and 2022 and a 94% probability of decline (Cleguer et al. 2023). Even in this well-established monitoring program, regional population surveys are typically conducted at intervals of approximately five years, limiting the capacity to resolve changes in distribution or occurrence between surveys. In contrast, the Arabian/Persian Gulf population is currently estimated at approximately 5,000 dugongs, including around 3,000 in United Arab Emirates (UAE) waters; however, historical reconstructions indicate an estimated 25% decline over the past 60 years (Al-Abdulrazzak & Pauly 2017). These regional trends emphasize a central challenge in dugong conservation: while monitoring is essential, repeated, large-scale population assessments are logistically demanding and often infrequent (Marsh et al. 2025a).

Dugong population assessments have traditionally relied on visual surveys from aircraft, and at smaller spatial scales, vessels and increasingly drone surveys (Alagha et al., 2025; Cleguer & Marsh, 2023). These approaches are essential as they can provide information on abundance, spatial distribution, group size and demographic composition. Despite their usefulness, visual surveys may be less effective for confirming presence at local scales where dugongs occur at low densities or intermittently. Environmental DNA (eDNA) offers a potentially valuable complementary approach for detecting dugong presence in discrete areas, particularly where animals are sparsely distributed or difficult to observe. When integrated with visual surveys, local knowledge or other monitoring approaches, eDNA detections could constitute an additional line of evidence for confirming dugong occurrence and identifying areas that need further investigation.

For rare and cryptic marine megafauna, eDNA analysis can provide sensitive presence data from water samples. Environmental DNA methods have been applied to detection of marine megafauna (Baker et al., 2018; Andruszkiewicz et al., 2017; Closek et al., 2019; Juhel et al., 2021; Qu & Stewart, 2019). For dugongs, Davis et al. (2026) used an eDNA metabarcoding approach to detect dugongs in the northern Saudi Red Sea, and highlighted the risk of false negatives in low-density populations. However, eDNA metabarcoding methods generally have lower limit of detection than single-species eDNA analysis, hereafter referred to as targeted eDNA analysis (Wood et al. 2019; Peng et al. 2023). Two other studies have developed targeted eDNA assays for dugong detection; yet, one lacked laboratory specificity testing and confirmatory sequencing (Bernagou et al. 2026), while the other used a SYBR-based assay and was only tested in the Saudi Arabian Red Sea coast (Mousavi-Derazmahalleh et al. 2026). These studies demonstrate the growing interest in dugong eDNA surveillance, but they also leave an important need for an assay that is analytically validated, confirmed under controlled conditions, tested across widely separated wild populations, and of high specificity. For this, probe-based TaqMan assays can offer greater confidence in species detection than SYBR-based approaches by reducing the risk of non-specific amplification and supporting future multiplexing (Charron et al. 2026)

Here, we developed and analytically validated a TaqMan qPCR assay for detecting dugong eDNA and evaluated its performance under controlled aquarium conditions and in wild populations from eastern Australia, north-western Australia and the Arabian/Persian Gulf. These regions encompass widely separated dugong populations and contrasting coastal environments, including turbid tropical Australian waters and the warm, high-irradiance waters of the Arabian/Persian Gulf. Our objectives were to: (1) develop a targeted eDNA assay and evaluate its specificity and analytical sensitivity against phylogenetically related and co-occurring species; (2) confirm detection of naturally shed dugong eDNA under controlled aquarium conditions; (3) validate field detections using confirmatory Sanger sequencing; and (4) assess the opportunities and limitations of targeted eDNA as a complementary surveillance tool across contrasting dugong habitats. By developing and validating a probe-based TaqMan eDNA assay across controlled and field settings in widely separated regions, with potential for future multiplexing, our study advances dugong eDNA research beyond assay development toward conservation-ready surveillance in marine environments.

## 2. MATERIALS AND METHODS

### 2.1 Environmental DNA assay development

We designed a species-specific TaqMan qPCR assay for dugong detection using mitochondrial genome sequences from the target species and seven co-occurring marine mammals downloaded from the National Center for Biotechnology Information (NCBI) database. The assay was developed independently of other recently published dugong-specific assays (Bernagou et al. 2026; Mousavi-Derazmahalleh et al. 2026). It is important to note that Mousavi-Derazmahalleh et al. (2026) developed a SYBR-based assay, which has potentially less specificity than TaqMan probe-based assays, as the latter allow incorporating additional mismatches with non-target species and permit multiplexing (Charron et al. 2026). Therefore, SYBR-based assays could need validation for the geographical region of interest (Charron et al. 2026).

Candidate primer and probe regions were identified from sequence alignments and screened *in-silico* using Primer-BLAST to assess specificity against non-target taxa (Supporting Information Table S1). Assay optimisation and specificity testing were conducted using tissue-derived DNA from dugongs and non-target species obtained from Queensland Museum (Australia), supplemented with turtle DNA extracts from species that share dugong habitats (described in Skinner et al. 2025). Tissue DNA was extracted, quantified, standardised, and used to evaluate primer performance, optimal annealing temperature, and amplification specificity.

The assay was optimised using temperature-gradient qPCR and subsequently tested for specificity against nine non-target marine vertebrates. The optimal annealing temperature was selected based on amplification performance, and assay sensitivity was assessed by estimating the limit of detection (LoD) and limit of quantification (loQ) using a serial dilution of synthetic double-stranded DNA fragments following Klymus et al. (2020). Detailed reaction compositions, cycling conditions, and analytical procedures are provided in the Supporting Information.

### 2.2 Environmental DNA sample collection

Water samples for assay validation were collected from two aquaria housing captive dugongs: SEA Life Sydney Aquarium (Australia) and SeaWorld Abu Dhabi (UAE). Field sampling included locations associated with recent dugong sightings or known dugong habitat from a total of eight coastal field sites in Cleveland Bay (north-east coast Australia), Broome (north-west coast Australia), and the Marawah Biosphere reserve in the Arabian/Persian Gulf (UAE). At each sampling location, between three and ten replicate water samples were filtered using self-preserving filter packs (5 µm MCE, Smith-Root), with aquarium and field blanks included to monitor contamination during sampling (Table 1, Supporting information Table S4).

**Table 1.**
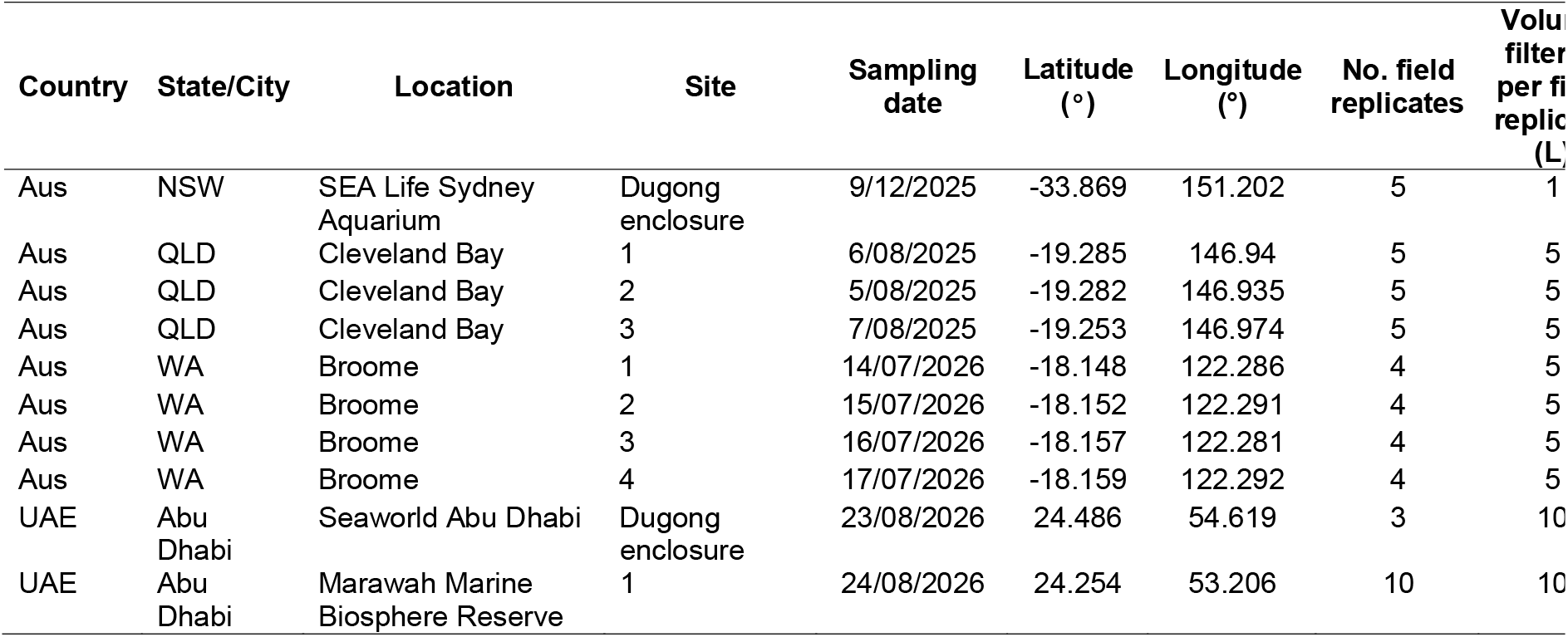
Information on aquaria and field sites where eDNA samples were collected for dugong eDNA detection using an assay targeting a 216 bp fragment of the *Dugong dugon* 16S mitochondrial gene. GPS coordinates recorded here belong to the first field replicate from each site. For a full list of GPS coordinates of each field replicate, please see Supporting information Table S4. Aus = Australia, UAE = United Arab Emirates. NSW = New South Wales, QLD = Queensland, and WA = Western Australia

### 2.3 Laboratory procedures

All eDNA sample analyses were carried out in the dedicated TropWATER eDNA Laboratory. Immediately upon return to the laboratory, filters were removed from filter packs and transferred to 2 mL DNA LoBind tubes containing 600 µL CTAB buffer and 60 µL proteinase K, then incubated overnight. The lysate was transferred to clean sterile 2 mL DNA LoBind tubes and centrifuged at 12,000 g for 10 min at ambient temperature. DNA from the supernatant was extracted using the Maxwell RSC PureFood GMO and Authentication Kit on a Maxwell RSC instrument following the manufacturer’s specifications. DNA was eluted in 100 µL UltraPure water and purified using the DNeasy PowerClean Pro Cleanup Kit (Qiagen). A negative extraction control was included with each batch of eDNA extractions to monitor contamination during laboratory procedures.

DNA extracts were tested for inhibition by spiking a known number of copies of a double-stranded synthetic DNA fragment from the armoured frog, *Litoria lorica*, into duplicate technical replicates from all field samples. The same number of copies was also spiked into three technical replicates containing laboratory-grade water. A sample was considered inhibited if a cycle threshold shift of three or more cycles was observed when comparing spiked field samples with spiked water controls.

All qPCR assays were run on a QuantStudio 5 Real-Time PCR System in white 384-well plates sealed with optical films. Environmental DNA detection was carried out using four technical replicates per aquarium/field sample. Each plate contained three no-template controls and a triplicated positive control consisting of synthetic double-stranded DNA fragments. Each qPCR reaction contained 5 µL TaqPath ProAmp Multiplex Master Mix, 0.5 µL forward primer (10 µM), 0.5 µL reverse primer (10 µM), 0.5 µL probe (10 µM), 0.5 µL UltraPure water, and 3 µL DNA extract. Thermal cycling conditions consisted of initial denaturation and activation at 95 °C for 10 min, followed by 40 cycles of 95 °C for 15 s, 60 °C for 1 min at a ramp rate of 1.6 °C/s, and 72 °C for 1 min at a ramp rate of 1.6 °C/s. Plates were analysed using the fluorescence threshold automatically set by QuantStudio Design and Analysis Software v.2.8.0 before export and analysis in Microsoft Excel. A subset of positive qPCR products was Sanger sequenced at the Australian Genome Research Facility in Brisbane to verify target-species amplification.

## 3. RESULTS

We developed an assay targeting a 216 bp fragment of the dugong 16S mitochondrial gene (Table 2), and *in-silico* and *in-vitro* analyses showed that the assay was specific to dugongs. Dugong eDNA was detected in all aquarium replicates from both countries. All Australian field replicates also showed positive amplification of dugong eDNA, and 30% of field replicates from the Arabian/Persian Gulf produced positive detections (Table 3). Sanger sequencing confirmed that all amplicons matched the dugong mitochondrial genome at ≥99% pairwise similarity (accession number NC_003314, positions 1,808–2,205).

**Table 2.**
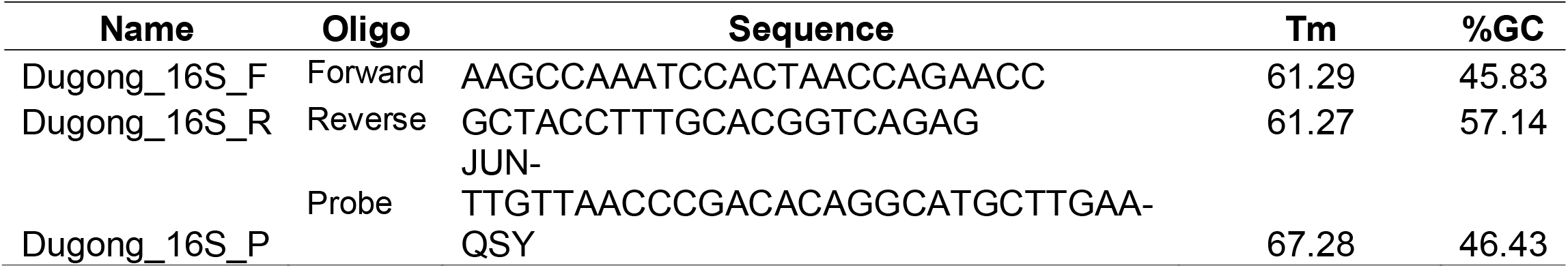
Information on primers and probe sequences of an assay targeting a 216 bp fragment of the *Dugong dugon* 16S mitochondrial gene.

**Table 3.** Environmental DNA detections of aquaria and field samples tested for dugong eDNA detection targeting a 216 bp fragment of the *gong dugon* 16S mitochondrial gene. Aus = Australia, UAE = United Arab Emirates. NSW = New South Wales, QLD = Queensland, and WA Western Australia. Red font indicates dugong eDNA detections in the corresponding site field blank

| Country | State/City | Location | Site | # field replicates | # positive field replicates | % positive field replicates | # qPCR technical replicates | # positive qPCR technical replicates | % positive qPCR technical replicates |
| --- | --- | --- | --- | --- | --- | --- | --- | --- | --- |
| Aus | NSW | SEA Life Sydney Aquarium | Dugong enclosure | 5 | 5 | 100 | 20 | 19 | 95 |
| Aus | QLD | Cleveland Bay | 1 | 5 | 5 | 100 | 20 | 10 | 50 |
| Aus | QLD | Cleveland Bay | 2 | 5 | 5 | 100 | 20 | 18 | 90 |
| Aus | QLD | Cleveland Bay | 3 | 5 | 5 | 100 | 20 | 14 | 70 |
| Aus | WA | Broome | 1 | 4 | 4 | 100 | 16 | 13 | 81.25 |
| Aus | WA | Broome | 2 | 4 | 4 | 100 | 16 | 15 | 93.75 |
| Aus | WA | Broome | 3 | 4 | 4 | 100 | 16 | 16 | 100 |
| Aus | WA | Broome | 4 | 4 | 4 | 100 | 16 | 6 | 37.5 |
| UAE | Abu Dhabi | Seaworld Abu Dhabi | Dugong enclosure | 3 | 3 | 100 | 12 | 12 | 100 |
| UAE | Abu Dhabi | Marawah Marine Biosphere Reserve | 1 | 10 | 3 | 30 | 40 | 3 | 7.5 |

## 4. DISCUSSION

Our results show that a validated, TaqMan probe-based assay can detect dugong eDNA across controlled and field settings spanning widely separated populations and contrasting marine environments, representing an internationally transferable tool for dugong surveillance. While other targeted eDNA assays for dugong detection have been recently published (Bernagou et al. 2026; Mousavi-Derazmahalleh et al. 2026), these have not been shown to work across geographically separated wild populations. Bernagou et al. (2026) did not include laboratory specificity of their assay testing or confirmatory sequencing of eDNA detections, while Mousavi-Derazmahalleh et al. (2026) developed a SYBR-based assay and only validated it for the Saudi Arabian Red Sea coast. The present study advances these approaches by using a probe-based TaqMan assay that offers an extra layer of specificity and evaluating its performance across geographically separated wild populations. Additionally, unlike SYBR-based assays, TaqMan assays allow for multiplexing, enabling end users to simultaneously screen for other co-occurring species of conservation concern, such as marine turtles. For conservation and management, this assay can help identify where dugongs have recently occurred, guide follow-up survey effort, and expand monitoring between infrequent broad-scale assessments.

The contrast of detectability between regions points to the need for improved understanding of detection probability when interpreting field non-detections. All field samples collected in Cleveland Bay and Broome produced positive detections. In contrast, a third of the field samples collected in the Marawah Marine Biosphere Reserve produced positive detections. These differences should not necessarily be interpreted as geographic differences in assay performance, especially because the aquarium samples from UAE resulted in 100% detections. The Australian sampling was undertaken in close association with field activities involving known dugong presence, including drone observations and dugong handling, and water was therefore collected in close proximity to animals. By contrast, samples in the UAE were collected after a herd of approximately 40–60 dugongs had been observed several hundred metres from the sampling location, with the animals subsequently moving through the area. Variable detection when dugongs are known to be present has been observed when using eDNA metabarcoding for dugong detection in the Red Sea despite samples being collected after dugong sightings (Davis et al. 2026). Similar limitations associated with interpreting eDNA non-detections have been recognised for other marine mammals and sirenians (Hunter et al. 2018; Suárez-Bregua et al. 2022).

Several biological and environmental processes could contribute to this variability. Environmental DNA in aquatic environments is subject to degradation, dilution and transport, with persistence and detectability influenced by factors including temperature, UV exposure, microbial activity and hydrodynamic processes (Strickler et al. 2015; Vorkapic et al. 2016; Barnes et al. 2021; Suárez-Bregua et al. 2022; Ballmer et al. 2024). In our study, water temperatures in the Marawah Marine Biosphere Reserve exceeded 32°C during sampling and the region experienced intense solar exposure, conditions that may favour more rapid eDNA degradation. Cleveland Bay and Broome are generally more turbid, potentially reducing UV penetration and photochemical degradation (Ballmer et al. 2024), although the extent to which this contributed to the observed difference is beyond the scope of this study.

Sampling design is also critical for eDNA detectability. Marine eDNA can be spatially heterogeneous, particularly for mobile animals occurring at relatively low density, making replication, sampling depth, spatial coverage and filtered water volume potentially important determinants of detection (Govindarajan et al. 2022; Suárez-Bregua et al. 2022). Also, collecting near-bottom water samples, consistent with the benthic foraging behaviour of dugongs, could improve detectability (Bernagou et al. 2026). Our results indicate that further optimisation should also experimentally evaluate timing and proximity of water sampling relative to animal presence, since the relative contribution of physicochemical and hydrodynamic processes cannot be separated from differences in animal proximity and sampling design in our dataset. The contrast between regions point to the need of an improved understanding of detection probability to interpret non-detections on field samples. Detection-probability modelling would help translate eDNA detections and non-detections into management-relevant inference.

Sample preservation and transport may have further influenced detection outcomes in the Arabian/Persian Gulf. Although self-preserving filters are intended to stabilise DNA after collection, prolonged exposure to elevated or fluctuating temperatures during transport may still reduce eDNA recovery. For operational dugong surveillance, especially in remote tropical regions, protocols should prioritise local processing, temperature-stable preservation, and rapid extraction where feasible.

The most immediate application of targeted dugong eDNA is likely to be local-scale presence surveillance, particularly in areas where dugongs occur at low density, use habitats intermittently, or are difficult to detect reliably using visual methods. Established aerial surveys remain essential for estimating dugong abundance and broad-scale distribution, while UAVs and other visual approaches can provide complementary information at finer spatial scales (Pollock et al. 2006; Cleguer & Marsh 2023; Alagha et al. 2025). These approaches provide ecological and demographic information that cannot currently be inferred reliably from eDNA. Targeted eDNA methods therefore have immediate value as a decision-support tool for dugong conservation. When embedded within standardised sampling designs and interpreted alongside visual surveys, local knowledge and habitat information, eDNA analysis can strengthen surveillance in areas where dugong presence is uncertain or difficult to confirm.

## Supporting information

Data archive following the FAIReDNA guidelines

## ACKNOWLEDGEMENTS

We thank Jessica Worthington Wilmer (Queensland Museum), Kerrie Dixon and Leia Cook (SEA Life Sydney Aquarium), and Elise Marquis and Chrissy Swad (SeaWorld Abu Dhabi). Luisa Schramm and Sarah Landeo Yauri collected Cleveland Bay samples. We gratefully acknowledge Nyamba Buru Yawuru and the Yawuru Traditional Owners, including the Yawuru hunters and rangers, for the Broome fieldwork. We thank Gysseth Herrera-Villarraga, Camilla Maniscalco, Oliane Pedersen, Sandra Infante-Villamil, John Kronenberger and Roy Barkan for laboratory analyses. We acknowledge the Environment Agency – Abu Dhabi for facilitating UAE field sampling. UAE samples were imported through the National Reference eDNA Centre, University of Canberra.

## SUPPORTING INFORMATION

Additional supporting information can be found online in the Supporting Information section of this article.

## DATA AVAILABILITY STATEMENT

Data is available and archived following the FAIR eDNA guidelines for accessible, available and reproducible data.

## REFERENCES

Al-Abdulrazzak, D., & Pauly, D. (2017). Reconstructing historical baselines for the Persian/Arabian Gulf dugong, Dugong dugon (Mammalia: Sirenia). Zoology in the Middle East, 63(2), 95–102. 10.1080/09397140.2017.1315853

Alagha, D. I., Soliman, Y., Marshall, C. D., & Alansi, M. (2025). Detection of the largest herd of dugongs (Dugong dugon) in the Central Arabian Gulf using unmanned aerial vehicles. Frontiers in Marine Science, 12, 1620194. 10.3389/fmars.2025.1620194

Andruszkiewicz, E. A., Starks, H. A., Chavez, F. P., Sassoubre, L. M., Block, B. A., & Boehm, A. B. (2017). Biomonitoring of marine vertebrates in Monterey Bay using eDNA metabarcoding. PLOS ONE, 12(4), e0176343. 10.1371/journal.pone.0176343

Baker, C. S., Steel, D., Nieukirk, S., & Klinck, H. (2018). Environmental DNA (eDNA) from the wake of the whales: Droplet digital PCR for detection and species identification. Frontiers in Marine Science, 5, 133. 10.3389/fmars.2018.00133

Ballmer, E., McNeill, K., & Deiner, K. (2024). Potential role of photochemistry in environmental DNA degradation. Environmental Science & Technology Letters, 11(12), 1284–1295. 10.1021/acs.estlett.4c00704

Barnes, M. A., Chadderton, W. L., Jerde, C. L., Mahon, A. R., Turner, C. R., & Lodge, D. M. (2021). Environmental conditions influence eDNA particle size distribution in aquatic systems. Environmental DNA, 3(3), 643–653. 10.1002/edn3.160

Bernagou, L., Serval, A., Beudard, F., Mathey, J., Mercky, Y., Le Bozec, C., Trotzuk, E., Paunde, K., Garcia, M., Bellemain, E., & Corse, E. (2026). Optimizing marine eDNA protocols for rare megafauna: A case study on dugongs in the Western Indian Ocean. Authorea. 10.22541/authorea.15003154/v1

Charron, M. R., Keber, B. M. L., Guxholli, R., Rogers, S., Ogden, J. L., Breault, P. W., & Docker, M. F. (2026). A multi-stage framework for validating targeted environmental DNA (eDNA) assays: From in silico design to in vitro and in situ testing for fisheries monitoring. Environmental Biology of Fishes. 10.1007/s10641-026-01868-y

Cleguer, C., Hamel, M., Rankin, R. W., Genson, A., Edwards, C., Collins, K., Crowe, M., Choukroun, S., & Marsh, H. (2023). 2022 dugong aerial survey: Mission Beach to Moreton Bay (JCU Centre for Tropical Water & Aquatic Ecosystem Research Publication No. 23/44). James Cook University. 10.25903/s661-1j55

Cleguer, C., & Marsh, H. (2023). An inventory of dugong aerial surveys in Australia (Report No. 23/22). TropWATER, James Cook University. https://www.nespmarinecoastal.edu.au/publication/1454/

Closek, C. J., Santora, J. A., Starks, H. A., Schroeder, I. D., Andruszkiewicz, E. A., Sakuma, K. M., Bograd, S. J., Hazen, E. L., Field, J. C., & Boehm, A. B. (2019). Marine vertebrate biodiversity and distribution within the Central California Current using environmental DNA (eDNA) metabarcoding and ecosystem surveys. Frontiers in Marine Science, 6. 10.3389/fmars.2019.00732

Davis, P. Z. R., Alghrair, M., Ramalho, R. O., & Poonian, C. N. S. (2026). Environmental DNA as an innovative tool to detect elusive dugong (Dugong dugon) populations. Aquatic Mammals, 52(4), 308–313. 10.1578/AM.52.4.2026.308

Govindarajan, A. F., McCartin, L., Adams, A., Allan, E., Belani, A., Francolini, R., Fujii, J., Gomez-Ibañez, D., Kukulya, A., Marin, F., Tradd, K., Yoerger, D. R., McDermott, J. M., & Herrera, S. (2022). Improved biodiversity detection using a large-volume environmental DNA sampler with in situ filtration and implications for marine eDNA sampling strategies. Deep Sea Research Part I: Oceanographic Research Papers, 189, 103871. 10.1016/j.dsr.2022.103871

He, Q., & Silliman, B. R. (2019). Climate change, human impacts, and coastal ecosystems in the Anthropocene. Current Biology, 29(19), R1021–R1035.

Hunter, M. E., Meigs-Friend, G., Ferrante, J., Dorazio, R., Keith-Diagne, L., Luna, F., Lanyon, J., & Reid, J. (2018). Surveys of environmental DNA (eDNA): A new approach to estimate occurrence in vulnerable manatee populations. Endangered Species Research, 35, 101–111. 10.3354/esr00880

Juhel, J., Marques, V., Polanco Fernández, A., Borrero-Pérez, G. H., Mutis Martinezguerra, M., Valentini, A., Dejean, T., Manel, S., Loiseau, N., Velez, L., Hocdé, R., Letessier, T. B., Richards, E., Hadjadj, F., Bessudo, S., Ladino, F., Albouy, C., Mouillot, D., & Pellissier, L. (2021). Detection of the elusive dwarf sperm whale (Kogia sima) using environmental DNA at Malpelo Island, Eastern Pacific, Colombia. Ecology and Evolution, 11(7), 2956–2962. 10.1002/ece3.7057

Klymus, K. E., Merkes, C. M., Allison, M. J., Goldberg, C. S., Helbing, C. C., Hunter, M. E., Jackson, C. A., Lance, R. F., Mangan, A. M., Monroe, E. M., Piaggio, A. J., Stokdyk, J. P., Wilson, C. C., & Richter, C. A. (2020). Reporting the limits of detection and quantification for environmental DNA assays. Environmental DNA, 2(3), 271–282. 10.1002/edn3.29

Marsh, H., Grech, A., & McMahon, K. (2018). Dugongs: Seagrass community specialists. In A. W. D. Larkum, G. A. Kendrick, & P. J. Ralph (Eds.), Seagrasses of Australia: Structure, ecology and conservation (pp. 629–661). Springer International Publishing. 10.1007/978-3-319-71354-0_19

Marsh, H., O’Shea, T. J., & Reynolds, J. E. (2011). Ecology and conservation of the Sirenia: Dugongs and manatees. Cambridge University Press.

Marsh, H., Alqahtani, T., Arosa, S., Beech, M., Blair, D., Burt, J. A., Das, H., Khamis, A., Loates, P., arshall, C., McKenzie, L., Pilcher, N., Rabaoui, L., & Wong, J. (2025a). Arabian/Persian Gulf. In H. Marsh, P. J. Loates, & L. Schramm (Eds.), Global assessment of dugong status and conservation needs (CMS Technical Series Publication No. 51, pp. 97–122). Convention on the Conservation of Migratory Species of Wild Animals. 10.25903/0ncx-v838

Marsh, H., Blair, D., Lafratta, A., Lanyon, J. M., & McKenzie, L. (2025b). Global context. In H. Marsh, P. J. Loates, & L. Schramm (Eds.), Global assessment of dugong status and conservation needs (CMS Technical Series Publication No. 51, pp. 1–43). Convention on the Conservation of Migratory Species of Wild Animals. 10.25903/0ncx-v838

Marsh, H., Loates, P. J., & Schramm, L. (2025c). A global assessment of dugong status and conservation needs (CMS Technical Series Publication No. 51). CMS Office – Abu Dhabi. 10.25903/0ncx-v838

McCauley, D. J., Pinsky, M. L., Palumbi, S. R., Estes, J. A., Joyce, F. H., & Warner, R. R. (2015). Marine defaunation: animal loss in the global ocean. Science, 347(6219), 1255641.

Mousavi-Derazmahalleh, M., Lukehurst, S. S., Hardenstine, R. S., Peverley, G., Dawkins, K. L., O’Toole, K. A., Asher, J. M., Boksmati, T., Richards, Z. T., Valsecchi, E., & Berumen, M. L. (2026). Development of a species-specific qPCR assay for the detection of the threatened dugong (Dugong dugon). Frontiers in Mammal Science, 5, 1915420. 10.3389/fmamm.2026.1915420

Peng, S., Wang, L., Ma, Y., Ye, L., Hou, C., Liu, Y., … & Dong, Z. (2023). Application of environmental DNA metabarcoding and quantitative PCR to detect blooming jellyfish in a temperate bay of northern China. Ecology and Evolution, 13(11), e10669.

Pimiento, C., Leprieur, F., Silvestro, D., Lefcheck, J. S., Albouy, C., Rasher, D. B., … & Griffin, J. N. (2020). Functional diversity of marine megafauna in the Anthropocene. Science Advances, 6(16), eaay7650.

Pollock, K. H., Marsh, H. D., Lawler, I. R., & Alldredge, M. W. (2006). Estimating animal abundance in heterogeneous environments: An application to aerial surveys for dugongs. Journal of Wildlife Management, 70(1), 255–262. 10.2193/0022-541X(2006)70[255:EAAIHE]2.0.CO;2

Qu, C., & Stewart, K. A. (2019). Evaluating monitoring options for conservation: Comparing traditional and environmental DNA tools for a critically endangered mammal. The Science of Nature, 106(3), 9. 10.1007/s00114-019-1605-1

Skinner, E., Hamann, M., Strugnell, J. M., & Rath, C. V. (2025). Investigating the limits of detection of environmental DNA from loggerhead and hawksbill turtles. Aquatic Conservation: Marine and Freshwater Ecosystems, 35(8), e70199. 10.1002/aqc.70199

Strickler, K. M., Fremier, A. K., & Goldberg, C. S. (2015). Quantifying effects of UV-B, temperature, and pH on eDNA degradation in aquatic microcosms. Biological Conservation, 183, 85–92. 10.1016/j.biocon.2014.11.038

Suárez-Bregua, P., Álvarez González, M., Parsons, K. M., Rotllant, J., Pierce, G. J., & Saavedra, C. (2022). Environmental DNA (eDNA) for monitoring marine mammals: Challenges and opportunities. Frontiers in Marine Science. 10.3389/fmars.2022.987774

Vorkapic, D., Pressler, K., & Schild, S. (2016). Multifaceted roles of extracellular DNA in bacterial physiology. Current Genetics, 62(1), 71–79. 10.1007/s00294-015-0514-x

Wirsing, A. J., Kiszka, J., Allen, A.-C., & Heithaus, M. R. (2022). Ecological roles and importance of sea cows (Order: Sirenia): A review and prospectus. Marine Ecology Progress Series, 689. 10.3354/meps14031

Wood, S. A., Pochon, X., Laroche, O., von Ammon, U., Adamson, J., & Zaiko, A. (2019). A comparison of droplet digital polymerase chain reaction (PCR), quantitative PCR and metabarcoding for species-specific detection in environmental DNA. Molecular Ecology Resources, 19(6), 1407–1419.

