## Supplementary information for "Finding dugongs through environmental DNA: a complementary tool for population surveillance"

**METHODS**

1. **Environmental DNA assay design and optimisation**

*In-silico* assay design consisted of downloading whole mitochondrial genome sequences of dugong (*Dugong dugon*) and seven co-occurring marine mammals from the National Center for Biotechnology Information database using Geneious Prime v.2025 2.2 (Table S1). Sequences were then aligned using the MUSCLE alignment algorithm (Edgar, 2004) to identify candidate primer and probe binding regions. Primer binding regions were selected based on the presence of mismatches at the 3’ end between *D. dugon* sequences and those of non-target species. *In-silico* specificity testing of candidate primers was carried out using Primer-BLAST (Ye et al., 2012).

Tissue samples for *in-vitro* assay optimisation and specificity testing were obtained from Queensland Museum, Australia (Table S2). Tissue DNA was extracted using the Cetyl Trimethylammonium Bromide protocol (Adamkewicz & Harasewych, 1996). DNA extracts from marine turtles described by Skinner et al. (2025) were also included in specificity testing because these species occupy the same habitats as dugongs (Table S2). DNA concentrations of target and non-target species were quantified on the Quantus instrument using the QuantiFluor ONE dsDNA System following the manufacturer’s protocol. All DNA extracts were diluted to concentrations of ~0.1 ng/µL and used as template for qPCR analyses.

Assay optimisation was performed using a temperature-gradient qPCR on a QuantStudio 3 Real-Time PCR System (Thermo Fisher Scientific), with dugong DNA extracts tested in triplicate. Each 10 µL reaction contained 5 µL SYBR Green Master Mix, 0.5 µL forward primer (10 µM), 0.5 µL reverse primer (10 µM), 1 µL UltraPure water, and 3 µL DNA template. Thermocycling conditions consisted of an initial denaturation at 95 °C for 10 min, followed by 40 cycles of 95 °C for 15 s and annealing at 58 °C, 60 °C, or 62 °C for 1 min, with a ramp rate of 1.6 °C/s. The optimal annealing temperature was selected as the temperature that produced the lowest quantification cycle values.

Assay specificity was then evaluated by testing DNA extracts from non-target species in triplicate on a QuantStudio 5 Real-Time PCR System (Thermo Fisher Scientific). Each 10 µL reaction contained 5 µL TaqPath ProAmp Multiplex Master Mix, 0.5 µL forward primer (10 µM), 0.5 µL reverse primer (10 µM), 0.5 µL probe (10 µM), 0.5 µL UltraPure water, and 3 µL DNA template. Thermocycling conditions followed those used for the temperature-gradient qPCR, using the selected optimal annealing temperature.

The assay’s limit of detection (LoD) and limit of quantification (LoQ) was determined using a 5-fold serial dilution of double-stranded synthetic DNA fragments (gBlocks, Integrated DNA Technologies) on a QuantStudio 5 Real-Time PCR System (Thermo Fisher Scientific). The dilution series ranged from 12500 DNA copies/reaction to 0.0032 DNA copies/reaction. qPCR wells were prepared using the same qPCR conditions applied for specificity testing and 12 qPCR technical replicates from each dilution. Limit-of-detection analysis was performed in RStudio v.2026.04.0 following Klymus et al. (2020).

**Table S1.** Accession numbers from marine mammal species downloaded from the National Center for Biotechnology Information (NCBI) using GeneiousPrime (v.2025 2.2) and used for in-silico primer and probe design

| **Species name** | **Accession no.** |
| --- | --- |
| *Dugong dugon* | NC_003314 |
| *Dugong dugon* | AJ421723 |
| *Dugong dugon* | AY075116 |
| *Dugong dugon* | PP134992 |
| *Stenella longirostris* | KX857351 |
| *Balaenoptera edeni* | NC_007938 |
| *Megaptera novaeangliae* | AP006467 |
| *Megaptera novaeangliae* | MF409246 |
| *Megaptera novaeangliae* | NC_006927 |
| *Megaptera novaeangliae* | PP475430 |
| *Orcaella heinsohni* | JF339977 |
| *Orcaella heinsohni* | NC_019591 |
| *Orcaella heinsohni* | PP829133 |
| *Orcaella heinsohni* | PP829132 |
| *Tursiops truncatus* | KT601198 |
| *Tursiops aduncus* | KF570335 |
| *Tursiops aduncus* | KF570336 |
| *Tursiops aduncus* | KF570337 |
| *Tursiops aduncus* | KF570338 |
| *Tursiops aduncus* | KF570339 |
| *Tursiops aduncus* | KF570340 |
| *Tursiops aduncus* | KF570341 |
| *Tursiops aduncus* | KF570344 |
| *Tursiops aduncus* | KF570343 |
| *Tursiops aduncus* | KF570342 |
| *Tursiops aduncus* | KF570359 |
| *Tursiops aduncus* | KF570360 |
| *Tursiops aduncus* | KF570361 |
| *Tursiops aduncus* | KF570362 |
| *Tursiops aduncus* | KF570353 |
| *Tursiops aduncus* | KF570354 |
| *Tursiops aduncus* | KF570355 |
| *Tursiops aduncus* | KF570356 |
| *Tursiops aduncus* | KF570357 |
| *Tursiops aduncus* | KF570358 |
| *Tursiops aduncus* | EU557092 |
| *Tursiops aduncus* | MG762973 |
| *Tursiops aduncus* | MG762974 |
| *Tursiops aduncus* | NC_012058 |
| *Sousa sahulensis* | KU522438 |

**Table S2.** Tissue samples used for *in-vitro* assay specificity testing

| **Sample origin** | **Sample no.** | **ID** | **Species name** | **Location** | **Latitude (°S)** | **Longitude (°E)** |
| --- | --- | --- | --- | --- | --- | --- |
| Queensland Museum | A013018 | D1 | *Dugong dugon* | Inskip Peninsula, between point and barge landing site | -25.826 | 153.042 |
| Queensland Museum | A014302 | D2 | *Dugong dugon* | Moore Park, Bundaberg | -24.719 | 152.287 |
| Queensland Museum | A014304 | D3 | *Dugong dugon* | Woodgate, Hervey Bay | -25.087 | 152.553 |
| Queensland Museum | A014307 | D4 | *Dugong dugon* | Mooroondu Park, Cleveland | -27.480 | 153.206 |
| Queensland Museum | A014308 | D5 | *Dugong dugon* | Four Mile Beach, Port Douglas | -16.508 | 145.467 |
| Queensland Museum | A014309 | D6 | *Dugong dugon* | Burnett Riverd | -24.759 | 152.390 |
| Queensland Museum | A014310 | D7 | *Dugong dugon* | Mackay | -20.636 | 148.712 |
| Queensland Museum | A014312 | D8 | *Dugong dugon* | Flinders Beach, North Stradbroke Island | -27.418 | 153.489 |
| Queensland Museum | A014313 | D9 | *Dugong dugon* | Port Clinton, Yeppoon | -23.500 | 150.717 |
| Queensland Museum | A014318 | D10 | *Dugong dugon* | Cleveland Bay, Townsville | -19.278 | 146.876 |
| Queensland Museum | A014319 | D11 | *Dugong dugon* | Peel Island | -27.526 | 153.335 |
| Queensland Museum | A014320 | D12 | *Dugong dugon* | Rainbow Channel, Moreton Bay | -27.419 | 153.413 |
| Queensland Museum | A014321 | D13 | *Dugong dugon* | Wellington Poiint | -27.469 | 153.267 |
| Queensland Museum | A014322 | D14 | *Dugong dugon* | Cape Grafton, Cairns | -16.863 | 145.917 |
| Queensland Museum | A014323 | D15 | *Dugong dugon* | Moore Park Beach, Bundaberg | -24.799 | 152.444 |
| Queensland Museum | A014326 | D16 | *Dugong dugon* | Clairview Beach | -22.118 | 149.536 |
| Queensland Museum | A014352 | D17 | *Dugong dugon* | Cooktown | -14.793 | 145.028 |
| Queensland Museum | A014980 | D18 | *Dugong dugon* | Yarrabah | -16.978 | 145.903 |
| Queensland Museum | A020509 | D19 | *Dugong dugon* | 4.1km N of Rainbow Beach | -20.846 | 153.075 |
| Queensland Museum | A014305 | M1 | *Megaptera*  *novaeangliae* | Blacks Beach, Mackay | -21.053 | 149.194 |
| Queensland Museum | A014306 | M2 | *Megaptera*  *novaeangliae* | Bribie Island | -27.061 | 153.202 |
| **Sample origin** | **Sample no.** | **ID** | **Species name** | **Location** | **Latitude (°S)** | **Longitude (°E)** |
| Queensland Museum | A014314 | M3 | *Megaptera*  *novaeangliae* | Mooloolaba River | -26.682 | 153.135 |
| Queensland Museum | A014325 | M4 | *Megaptera*  *novaeangliae* | Off Cowan Point, Moreton Island | -27.145 | 153.329 |
| Queensland Museum | A014327 | M5 | *Megaptera*  *novaeangliae* | Off Caloundra/Bribie Island | -26.812 | 153.137 |
| Queensland Museum | A014337 | M6 | *Megaptera*  *novaeangliae* | Moreton Island | -27.104 | 153.443 |
| Queensland Museum | A014338 | M7 | *Megaptera*  *novaeangliae* | Whitsunday Islands | -20.161 | 148.823 |
| Queensland Museum | A014339 | M8 | *Megaptera*  *novaeangliae* | Fraser Island | -25.525 | 153.122 |
| Queensland Museum | A015023 | M9 | *Megaptera*  *novaeangliae* | North Stradbroke Island | -27.518 | 153.505 |
| Queensland Museum | A015038 | M10 | *Megaptera*  *novaeangliae* | Warana Beach, Bokarina, Sunshine Coast | -26.741 | 153.136 |
| Queensland Museum | A015928 | M11 | *Megaptera*  *novaeangliae* | Main Beach, Gold Coast | -27.976 | 153.430 |
| Queensland Museum | A015930 | M12 | *Megaptera*  *novaeangliae* | Peregian Beach (south) | -26.501 | 153.095 |
| Queensland Museum | A023778 | M13 | *Megaptera*  *novaeangliae* | via Tewantin | -26.383 | 153.017 |
| Queensland Museum | A007770 | O1 | *Orcaella heinsohni* | Two-Mile Creek Toolakea Beach | -19.128 | 146.545 |
| Queensland Museum | A007771 | O2 | *Orcaella heinsohni* | Two-Mile Creek Toolakea Beach | -19.128 | 146.545 |
| Queensland Museum | A014311 | O3 | *Orcaella heinsohni* | Cooee Bay, Yeppoon | -23.143 | 150.761 |
| Queensland Museum | A003253 | S1 | *Sousa sahulensis* | Mouth of Brisbane River | -27.364 | 153.154 |
| Queensland Museum | A015933 | S2 | *Sousa sahulensis* | Gold Coast | -28.143 | 153.507 |
| Queensland Museum | A015937 | S3 | *Sousa sahulensis* | Deception Bay | -27.195 | 153.036 |
| Queensland Museum | A015932 | T4 | Tursiops (unidentified) | Eastern Beach, Fraser Island | -25.682 | 153.077 |
| Queensland Museum | A015934 | T5 | Tursiops (unidentified) | Kings Beach, Caloundra | -26.808 | 153.146 |
| Queensland Museum | A014328 | T1 | *Tursiops aduncus* | Cape Bedford Beach | -15.307 | 145.294 |
| Queensland Museum | A014340 | T2 | *Tursiops aduncus* | Lammermoor Beach | -23.153 | 150.767 |
| Queensland Museum | A015929 | T3 | *Tursiops aduncus* | Macdonald Point, Shoalwater Bay | -22.649 | 150.789 |
| **Sample origin** | **Sample no.** | **ID** | **Species name** | **Location** | **Latitude (°S)** | **Longitude (°E)** |
| Skinner et al. 2025 | - | CC1 | *Caretta caretta* |  |  |  |
| Skinner et al. 2025 | - | CC2 | *Caretta caretta* |  |  |  |
| Skinner et al. 2025 | - | CC3 | *Caretta caretta* |  |  |  |
| Skinner et al. 2025 | - | A8 | *Chelonia mydas* |  |  |  |
| Skinner et al. 2025 | - | A15 | *Chelonia mydas* |  |  |  |
| Skinner et al. 2025 | - | A20 | *Chelonia mydas* |  |  |  |
| Skinner et al. 2025 | - | FB | *Natator depressus* |  |  |  |
| Skinner et al. 2025 | - | EI1 | *Eretmochelys imbricata* |  |  |  |
| Skinner et al. 2025 | - | EI2 | *Eretmochelys imbricata* |  |  |  |

### RESULTS

1. **Environmental DNA assay design and optimisation**

The temperature gradient qPCR revealed marginal differences in Cq values across the three tested annealing temperatures. We chose 60°C as optimal annealing temperature given that it is commonly used across other eDNA assays, allowing for multiplexing in the future (Table S3). Specificity testing revealed that the candidate assay only amplified the genomic DNA of *D. dugon*.

**Table S3.** Results of the temperature gradient qPCR performed to determine the optimal annealing temperature of the candidate eDNA primers

| **Annealing temperature (°C)** | **Standard** | **Number of replicates** | **Cq Mean** | **Cq SD** |
| --- | --- | --- | --- | --- |
| 58 | Std1 | 3 | 17.3909 | 0.3207 |
|  | Std2 | 3 | 20.8784 | 0.2242 |
|  | Std3 | 3 | 24.7369 | 0.2795 |
| 60 | Std1 | 3 | 17.3084 | 0.3065 |
|  | Std2 | 3 | 20.8023 | 0.2265 |
|  | Std3 | 3 | 24.6046 | 0.2911 |
| 62 | Std1 | 3 | 17.4327 | 0.2809 |
|  | Std2 | 3 | 20.9454 | 0.2372 |
|  | Std3 | 3 | 24.7242 | 0.3281 |

The assay’s LoD and LoQ were estimated to be 1.3 DNA copies/reaction and 44 DNA copies/reaction, respectively (efficiency = 103.37% and R^2^ = 0.992) (Figure S1).

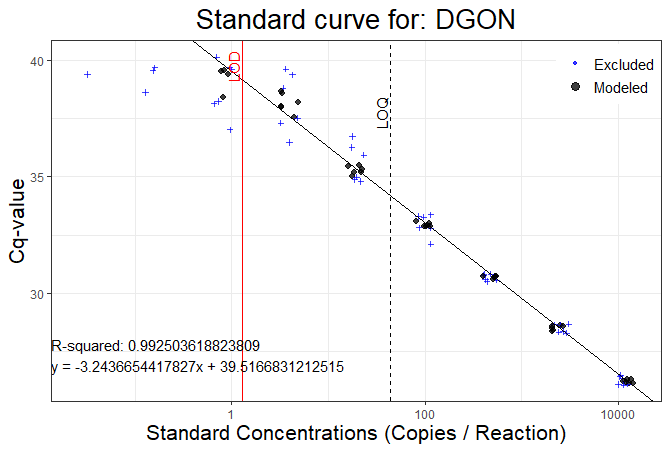

**Figure S1.** Modelled limit of detection (LoD, vertical red line) and limit of quantification (LoQ, vertical dashed line) of an assay targeting a 216 bp fragment of the *Dugong dugon* mitochondrial genome

2. **Environmental DNA sampling**

Experimental validation of the eDNA assay consisted of filtering water from aquaria housing individual dugongs in Sydney (SEA Life Sydney Aquarium) and Abu Dhabi (SeaWorld Abu Dhabi). Between three and five replicate water samples and one aquarium blank were filtered using self-preserving filter packs (5 µm MCE, Smith-Root) at each facility.

Field validation consisted of collecting eDNA samples at three separate sites on the east coast of Australia, four sites on the northwest coast of Australia and one site on the Marawah Marine Biosphere Reserve in the United Arab Emirates. At each site, between four and ten field replicates were collected and 5 – 10 L of water was filtered using self-preserving filter packs (5 µm MCE, Smith-Root). A field blank was also included at each site to monitor for field-induced contamination. A detailed list of the field samples is given in Table S4.

**Table S4.** Information on aquaria and field sites where eDNA samples were collected for dugong eDNA detection using an assay targeting a 216 bp fragment of the *Dugong dugon* 16S mitochondrial gene

| **Country** | **State/City** | **Location** | **Site** | **Sample ID** | **Sampling date** | **Latitude (°)** | **Longitude (°)** | **Filter pore size (µm)** | **Volume fitlered (L)** |
| --- | --- | --- | --- | --- | --- | --- | --- | --- | --- |
| Australia | New South Wales | SEA Life Sydney Aquarium | Dugong enclosure | Sea Life rep1 | 9/12/2025 | -33.869 | 151.202 | 5 | 1 |
| Australia | New South Wales | SEA Life Sydney Aquarium | Dugong enclosure | Sea Life rep2 | 9/12/2025 | -33.869 | 151.202 | 5 | 1 |
| Australia | New South Wales | SEA Life Sydney Aquarium | Dugong enclosure | Sea Life rep3 | 9/12/2025 | -33.869 | 151.202 | 5 | 1 |
| Australia | New South Wales | SEA Life Sydney Aquarium | Dugong enclosure | Sea Life rep4 | 9/12/2025 | -33.869 | 151.202 | 5 | 1 |
| Australia | New South Wales | SEA Life Sydney Aquarium | Dugong enclosure | Sea Life rep5 | 9/12/2025 | -33.869 | 151.202 | 5 | 1 |
| Australia | New South Wales | SEA Life Sydney Aquarium | Dugong enclosure | Sea Life Control | 9/12/2025 | -33.869 | 151.202 | 5 | 1 |
| Australia | Queensland | Cleveland Bay | 1 | Site 1 rep1 | 6/08/2025 | -19.285 | 146.940 | 5 | 5 |
| Australia | Queensland | Cleveland Bay | 1 | Site 1 rep2 | 6/08/2025 | -19.285 | 146.940 | 5 | 5 |
| Australia | Queensland | Cleveland Bay | 1 | Site 1 rep3 | 6/08/2025 | -19.284 | 146.940 | 5 | 5 |
| Australia | Queensland | Cleveland Bay | 1 | Site 1 rep4 | 6/08/2025 | -19.284 | 146.940 | 5 | 5 |
| Australia | Queensland | Cleveland Bay | 1 | Site 1 rep5 | 6/08/2025 | -19.282 | 146.940 | 5 | 5 |
| Australia | Queensland | Cleveland Bay | 1 | Site 1 Control | 6/08/2025 | -19.285 | 146.940 | 5 | 1 |
| Australia | Queensland | Cleveland Bay | 2 | Site 2 rep1 | 5/08/2025 | -19.282 | 146.935 | 5 | 5 |
| Australia | Queensland | Cleveland Bay | 2 | Site 2 rep2 | 5/08/2025 | -19.281 | 146.935 | 5 | 5 |
| Australia | Queensland | Cleveland Bay | 2 | Site 2 rep3 | 5/08/2025 | -19.283 | 146.935 | 5 | 5 |
| Australia | Queensland | Cleveland Bay | 2 | Site 2 rep4 | 5/08/2025 | -19.283 | 146.935 | 5 | 5 |
| Australia | Queensland | Cleveland Bay | 2 | Site 2 rep5 | 5/08/2025 | -19.283 | 146.935 | 5 | 5 |
| Australia | Queensland | Cleveland Bay | 2 | Site 2 Control | 5/08/2025 | -19.282 | 146.935 | 5 | 1 |
| **Country** | **State/City** | **Location** | **Site** | **Sample ID** | **Sampling date** | **Latitude (°)** | **Longitude (°)** | **Filter pore size (µm)** | **Volume fitlered (L)** |
| Australia | Queensland | Cleveland Bay | 3 | Site 3 rep1 | 7/08/2025 | -19.253 | 146.974 | 5 | 5 |
| Australia | Queensland | Cleveland Bay | 3 | Site 3 rep2 | 7/08/2025 | -19.251 | 146.974 | 5 | 5 |
| Australia | Queensland | Cleveland Bay | 3 | Site 3 rep3 | 7/08/2025 | -19.253 | 146.968 | 5 | 5 |
| Australia | Queensland | Cleveland Bay | 3 | Site 3 rep4 | 7/08/2025 | -19.252 | 146.968 | 5 | 5 |
| Australia | Queensland | Cleveland Bay | 3 | Site 3 rep5 | 7/08/2025 | -19.251 | 146.968 | 5 | 5 |
| Australia | Queensland | Cleveland Bay | 3 | Site 3 Control | 7/08/2025 | -19.253 | 146.974 | 5 | 1 |
| Australia | Western Australia | Broome | 1 | Site 1 rep1 | 14/07/2026 | -18.148 | 122.286 | 5 | 5 |
| Australia | Western Australia | Broome | 1 | Site 1 rep2 | 14/07/2026 | -18.148 | 122.286 | 5 | 5 |
| Australia | Western Australia | Broome | 1 | Site 1 rep3 | 14/07/2026 | -18.148 | 122.286 | 5 | 5 |
| Australia | Western Australia | Broome | 1 | Site 1 rep4 | 14/07/2026 | -18.148 | 122.286 | 5 | 5 |
| Australia | Western Australia | Broome | 1 | Site 1 Control | 14/07/2026 | -18.148 | 122.286 | 5 | 1 |
| Australia | Western Australia | Broome | 2 | Site 2 rep1 | 15/07/2026 | -18.152 | 122.291 | 5 | 5 |
| Australia | Western Australia | Broome | 2 | Site 2 rep2 | 15/07/2026 | -18.152 | 122.291 | 5 | 5 |
| Australia | Western Australia | Broome | 2 | Site 2 rep3 | 15/07/2026 | -18.152 | 122.291 | 5 | 5 |
| Australia | Western Australia | Broome | 2 | Site 2 rep4 | 15/07/2026 | -18.152 | 122.291 | 5 | 5 |
| Australia | Western Australia | Broome | 2 | Site 2 Control | 15/07/2026 | -18.152 | 122.291 | 5 | 1 |
| **Country** | **State/City** | **Location** | **Site** | **Sample ID** | **Sampling date** | **Latitude (°)** | **Longitude (°)** | **Filter pore size (µm)** | **Volume fitlered (L)** |
| Australia | Western Australia | Broome | 3 | Site 3 rep1 | 16/07/2026 | -18.157 | 122.281 | 5 | 5 |
| Australia | Western Australia | Broome | 3 | Site 3 rep2 | 16/07/2026 | -18.157 | 122.281 | 5 | 5 |
| Australia | Western Australia | Broome | 3 | Site 3 rep3 | 16/07/2026 | -18.157 | 122.281 | 5 | 5 |
| Australia | Western Australia | Broome | 3 | Site 3 rep4 | 16/07/2026 | -18.157 | 122.281 | 5 | 5 |
| Australia | Western Australia | Broome | 3 | Site 3 Control | 16/07/2026 | -18.157 | 122.281 | 5 | 1 |
| Australia | Western Australia | Broome | 4 | Site 4 rep1 | 17/07/2026 | -18.159 | 122.292 | 5 | 5 |
| Australia | Western Australia | Broome | 4 | Site 4 rep2 | 17/07/2026 | -18.159 | 122.292 | 5 | 5 |
| Australia | Western Australia | Broome | 4 | Site 4 rep3 | 17/07/2026 | -18.159 | 122.292 | 5 | 5 |
| Australia | Western Australia | Broome | 4 | Site 4 rep4 | 17/07/2026 | -18.159 | 122.292 | 5 | 5 |
| Australia | Western Australia | Broome | 4 | Site 4 Control | 17/07/2026 | -18.159 | 122.292 | 5 | 1 |
| United Arab Emirates | Abu Dhabi | Seaworld Abu Dhabi | Dugong enclosure | Sea World 1 | 23/08/2026 | 24.486 | 54.619 | 5 | 10 |
| United Arab Emirates | Abu Dhabi | Seaworld Abu Dhabi | Dugong enclosure | Sea World 2 | 23/08/2026 | 24.486 | 54.619 | 5 | 10 |
| United Arab Emirates | Abu Dhabi | Seaworld Abu Dhabi | Dugong enclosure | Sea World 3 | 23/08/2026 | 24.486 | 54.619 | 5 | 10 |
| **Country** | **State/City** | **Location** | **Site** | **Sample ID** | **Sampling date** | **Latitude (°)** | **Longitude (°)** | **Filter pore size (µm)** | **Volume fitlered (L)** |
| United Arab Emirates | Abu Dhabi | Seaworld Abu Dhabi | Dugong enclosure | Sea World Control | 23/08/2026 | 24.486 | 54.619 | 5 | 2 |
| United Arab Emirates | Abu Dhabi | Marawah Marine Biosphere Reserve | 1 | Site 1 rep1 | 24/08/2026 | 24.254 | 53.206 | 5 | 10 |
| United Arab Emirates | Abu Dhabi | Marawah Marine Biosphere Reserve | 1 | Site 1 rep4 | 24/08/2026 | 24.254 | 53.206 | 5 | 10 |
| United Arab Emirates | Abu Dhabi | Marawah Marine Biosphere Reserve | 1 | Site 1 rep9 | 24/08/2026 | 24.254 | 53.206 | 5 | 10 |
| United Arab Emirates | Abu Dhabi | Marawah Marine Biosphere Reserve | 1 | Site 1 rep2 | 24/08/2026 | 24.254 | 53.205 | 5 | 10 |
| United Arab Emirates | Abu Dhabi | Marawah Marine Biosphere Reserve | 1 | Site 1 rep3 | 24/08/2026 | 24.254 | 53.205 | 5 | 10 |
| United Arab Emirates | Abu Dhabi | Marawah Marine Biosphere Reserve | 1 | Site 1 rep5 | 24/08/2026 | 24.254 | 53.205 | 5 | 10 |
| United Arab Emirates | Abu Dhabi | Marawah Marine Biosphere Reserve | 1 | Site 1 rep6 | 24/08/2026 | 24.254 | 53.205 | 5 | 10 |
| United Arab Emirates | Abu Dhabi | Marawah Marine Biosphere Reserve | 1 | Site 1 rep7 | 24/08/2026 | 24.254 | 53.205 | 5 | 10 |
| United Arab Emirates | Abu Dhabi | Marawah Marine Biosphere Reserve | 1 | Site 1 rep8 | 24/08/2026 | 24.254 | 53.205 | 5 | 10 |
| United Arab Emirates | Abu Dhabi | Marawah Marine Biosphere Reserve | 1 | Site 1 rep10 | 24/08/2026 | 24.254 | 53.205 | 5 | 10 |
| United Arab Emirates | Abu Dhabi | Marawah Marine Biosphere Reserve | 1 | Site 1 Control | 24/08/2026 | 24.254 | 53.205 | 5 | 2 |

### References

Adamkewicz, S. L., & Harasewych, M. (1996). *Systematics and biogeography of the genus Donax (Bivalvia: Donacidae) in Eastern North America*. American Malacological Bulletin, 13, 97–103.

Edgar, R. C. (2004). MUSCLE: Multiple sequence alignment with high accuracy and high throughput. *Nucleic Acids Research, 32*(5), 1792–1797. <https://doi.org/10.1093/nar/gkh340>

Klymus, K. E., Merkes, C. M., Allison, M. J., Goldberg, C. S., Helbing, C. C., Hunter, M. E., Jackson, C. A., Lance, R. F., Mangan, A. M., Monroe, E. M., Piaggio, A. J., Stokdyk, J. P., Wilson, C. C., & Richter, C. A. (2020). Reporting the limits of detection and quantification for environmental DNA assays. *Environmental DNA, 2*(3), 271–282. <https://doi.org/10.1002/edn3.29>

Skinner, E., Hamann, M., Strugnell, J. M., & Rath, C. V. (2025). Investigating the Limits of Detection of Environmental DNA From Loggerhead and Hawksbill Turtles. *Aquatic Conservation: Marine and Freshwater Ecosystems, 35*(8), e70199. <https://doi.org/10.1002/aqc.70199>

Ye, J., Coulouris, G., Zaretskaya, I., Cutcutache, I., Rozen, S., & Madden, T. L. (2012). Primer-BLAST: A tool to design target-specific primers for polymerase chain reaction. *BMC Bioinformatics, 13*(1), 134. <https://doi.org/10.1186/1471-2105-13-134>
